# Essential Resources for Curated Immune Markers and Antibody Panels in Cytometry Analysis

**DOI:** 10.1101/2025.10.13.682059

**Authors:** Seongyun Choi, Jaeho Ji, Eunjeong Han, Ahrum Son, Hyunsoo Kim

## Abstract

The immune system, fundamental to numerous biological research fields, is increasingly investigated at single-cell resolution using high-dimensional flow cytometry and mass cytometry technologies. These approaches require comprehensive immunomarker information and carefully designed antibody panels that address the technical challenges of isotopic and spectral overlap inherent in metal/fluorescent-labeled antibodies. While immune cell markers have been extensively characterized and systematically catalogued, antibody panel information remains fragmented across the literature. To bridge this critical gap, we present ImapDB (Immune cell Marker and Panel Database), a novel Python-based graphical user interface that consolidates curated information on cell markers and antibody panels. ImapDB integrates data from commercial guidebooks, peer-reviewed literature, and validated research sources, currently focusing on normal human immune cell states. The database encompasses 127 distinct immune cell subtypes with over 350 validated markers and includes 45 commercial and 23 custom research antibody panels. Available on GitHub (https://github.com/kimlab-cnu/Cell_Marker_GUI), ImapDB serves as a centralized repository designed to streamline immune profiling assays and enhance research efficiency.

## Introduction

Cytology has evolved from traditional microscopic examination of individual cells to encompass sophisticated single-cell analysis technologies across diverse research domains including pathology, oncology, proteomics, and transcriptomics [1, 2]. These technological advances have been particularly transformative in immunology, enabling unprecedented insights into immune system complexity. Recent applications of single-cell approaches have revealed novel immune cell subtypes in cancer immunotherapy responses, such as the identification of exhausted CD8+ T cell populations that predict checkpoint inhibitor efficacy, and have uncovered previously unknown developmental trajectories in autoimmune diseases including the discovery of pathogenic T helper cell populations in multiple sclerosis [3].

Central to these advances are flow cytometry (FACS, Fluorescence-Activated Cell Sorting) and mass cytometry (CyTOF, Cytometry by Time-Of-Flight), which enable precise characterization of immune cell populations through multi-parametric analysis capable of simultaneously measuring up to 50 parameters per cell [3]. Immune cells, broadly categorized as myeloid cells and lymphocytes, express distinctive protein markers on their surface, intracellularly, and as secreted factors [4]. These markers serve as molecular signatures for cell identity; for instance, CD3 defines T lymphocytes, CD19 identifies B cells, CD14 marks classical monocytes, and CD56 characterizes natural killer cells. The expression patterns of these markers dynamically change throughout cell differentiation, activation states, and in response to environmental stimuli, making their systematic organization essential for accurate immunological studies [5].

Multi-marker analysis requires carefully designed antibody panels or cocktails comprising multiple antibodies that can simultaneously detect various cellular markers. The interpretation of co-expression patterns provides crucial insights into cell identity and functional states. For example, the combination of CD3+CD4+CD25+CD127low identifies regulatory T cells, while CD3+CD8+PD-1+TIM-3+ marks exhausted cytotoxic T cells [6-8]. However, panel design presents significant technical challenges, particularly in managing interference effects between metal isotopes in mass cytometry, where signal spillover can exceed 2% between adjacent channels, or fluorescence spectral overlap in flow cytometry, where compensation requirements can limit the number of usable fluorophore combinations [9, 10].

Current approaches to panel design rely predominantly on commercial tools with limited flexibility or institutional services that may not accommodate specialized research needs. While commercial panels offer standardized solutions, researchers frequently require custom configurations tailored to specific biological questions. The design process demands comprehensive knowledge of marker expression patterns, antibody performance characteristics, and platform-specific technical considerations [11]. Despite the critical importance of panel optimization, no comprehensive resource currently integrates marker information with validated antibody configurations.

We developed ImapDB to address this gap by providing a unified platform that combines systematic marker organization with curated antibody panel information. This Python-based graphical user interface consolidates data from authoritative sources, offering researchers immediate access to validated protocols and enabling informed experimental design. By bridging the gap between theoretical marker knowledge and practical implementation, ImapDB facilitates standardized approaches to immune profiling while supporting innovative applications in immunological research.

## Materials and Methods

### Software Architecture and Development

ImapDB was developed as a Python-based application utilizing PyQt5 for graphical interface implementation and Qt Designer for interface layout. The software architecture employs a modular design with separate components for data management, user interface, and file input/output operations. Data processing capabilities were implemented using Pandas for efficient manipulation of tabular data, while openpyxl provides compatibility with Excel formats commonly used in laboratory workflows **(Figure 1)**.

**Figure 1.**
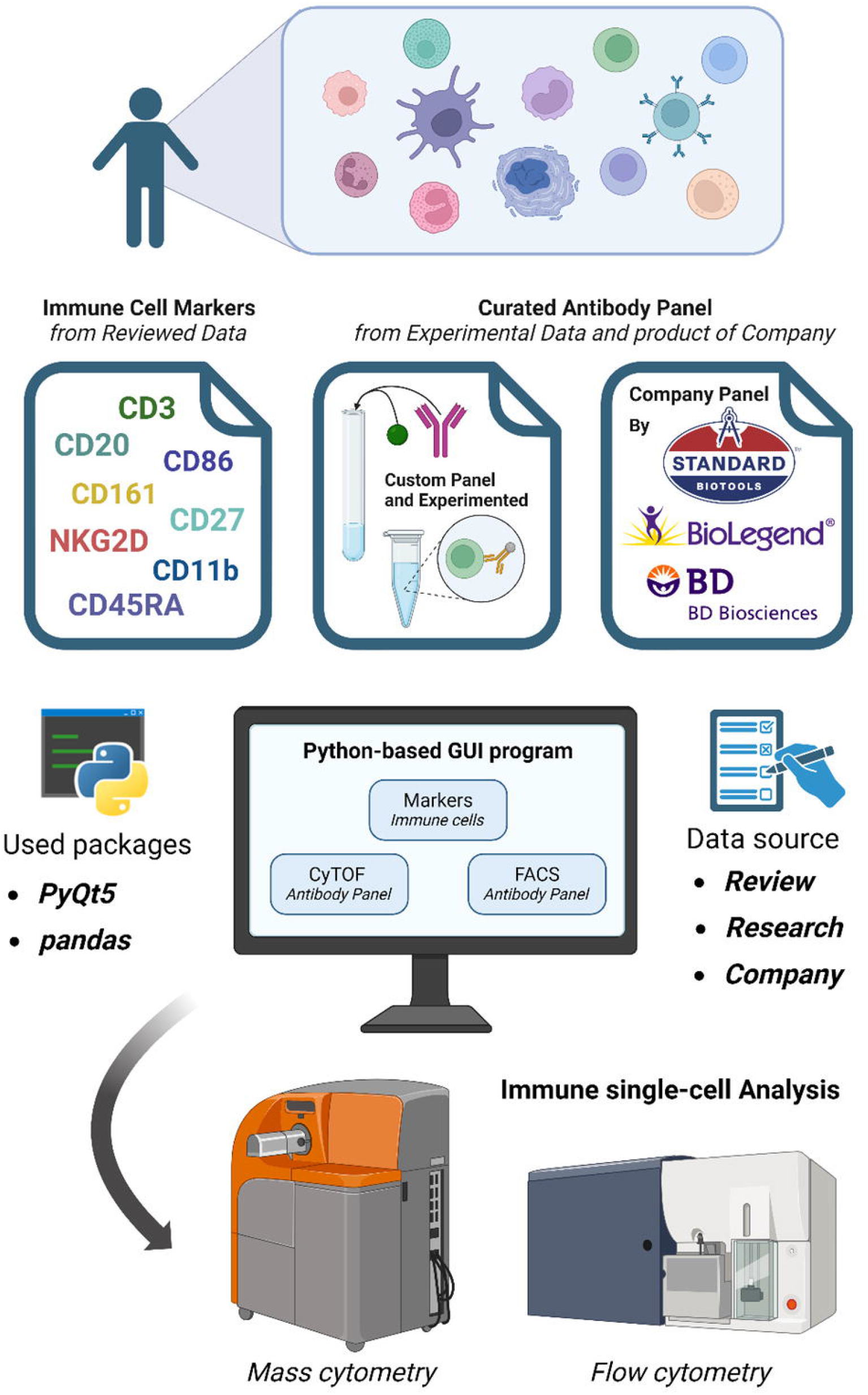
Overview of ImapDB architecture and workflow. The database integrates immune cell marker information with antibody panel configurations through a Python-based graphical user interface, providing hierarchical navigation and comprehensive search capabilities.

### System Requirements and Installation

The application operates on Windows, macOS, and Linux platforms with Python 3.6 or higher. Required dependencies include PyQt5 (installation via pip install pyqt5 and pip install pyqt5-tools), Pandas (pip install pandas), and openpyxl (pip install openpyxl). Following dependency installation, users can download the complete package from our GitHub repository. Path configuration may be required to ensure proper file loading, with detailed instructions provided in the documentation.

### Database Structure and Navigation

ImapDB implements a four-module architecture comprising a main navigation window and three specialized interfaces for immune markers, CyTOF panels, and FACS panels **(Figure 2)**. The hierarchical organization facilitates intuitive navigation through cascading selection menus that reflect biological relationships between cell populations. The Immune Dialog enables marker searches through three-tier selection: lineage (Lymphocyte/Myeloid), cell type, and subpopulation. Panel interfaces distinguish between commercial and experimental resources, each accessible through dedicated selection buttons **(Figure 3)**.

**Figure 2.**
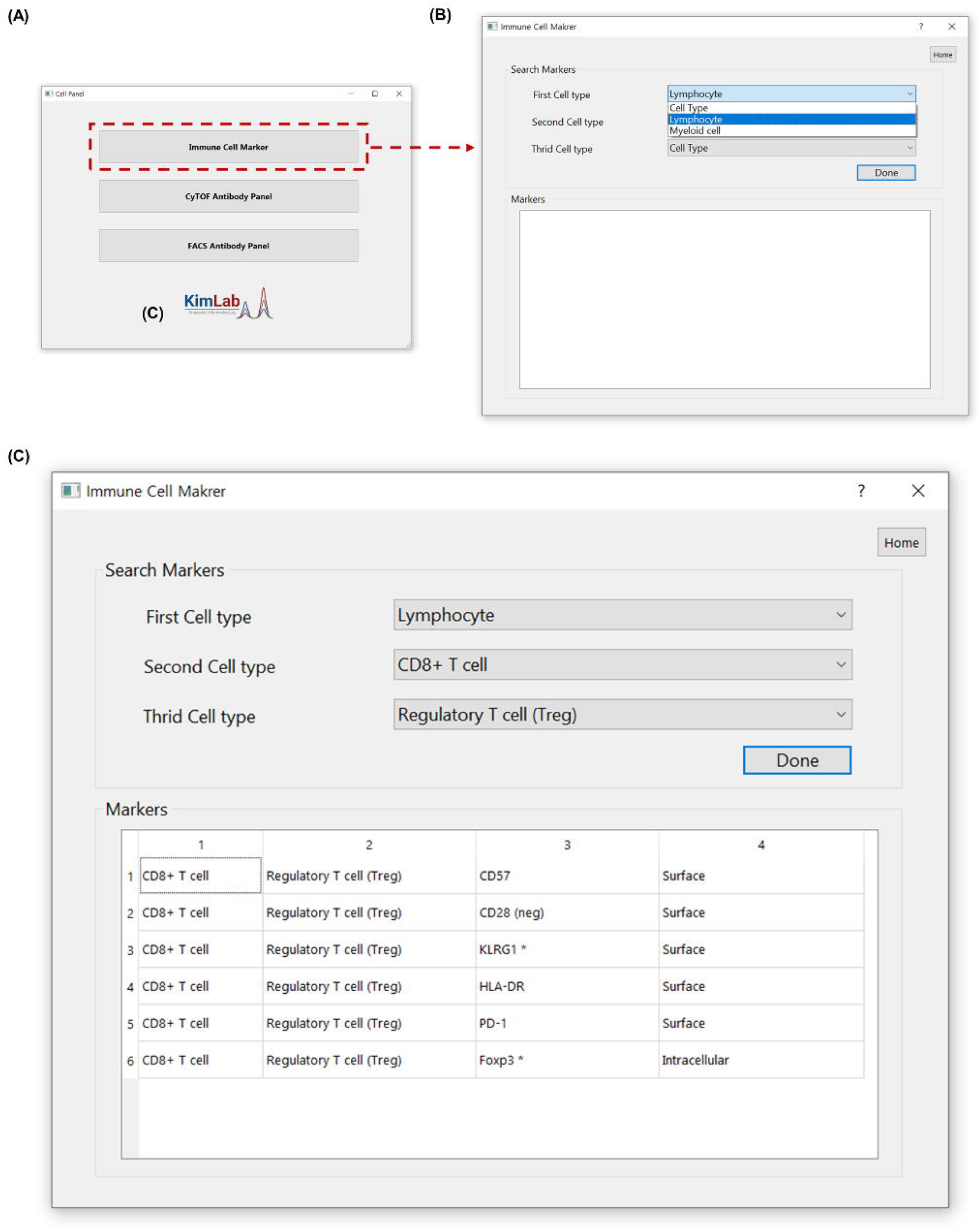
Hierarchical navigation interface for immune cell marker identification in ImapDB. (A) Main application window displaying three functional modules for accessing immune cell markers, CyTOF antibody panels, and FACS antibody panels. (B) Navigation pathway from the main window to the Immune Cell Marker Dialog through button selection. (C) Three-tier hierarchical selection system demonstrating marker identification for regulatory T cells, with sequential selection of cell lineage (Lymphocyte), cell type (CD4+ T cell), and subpopulation (Regulatory T cell). The resulting marker profile is displayed in tabular format below the selection interface.

**Figure 3.**
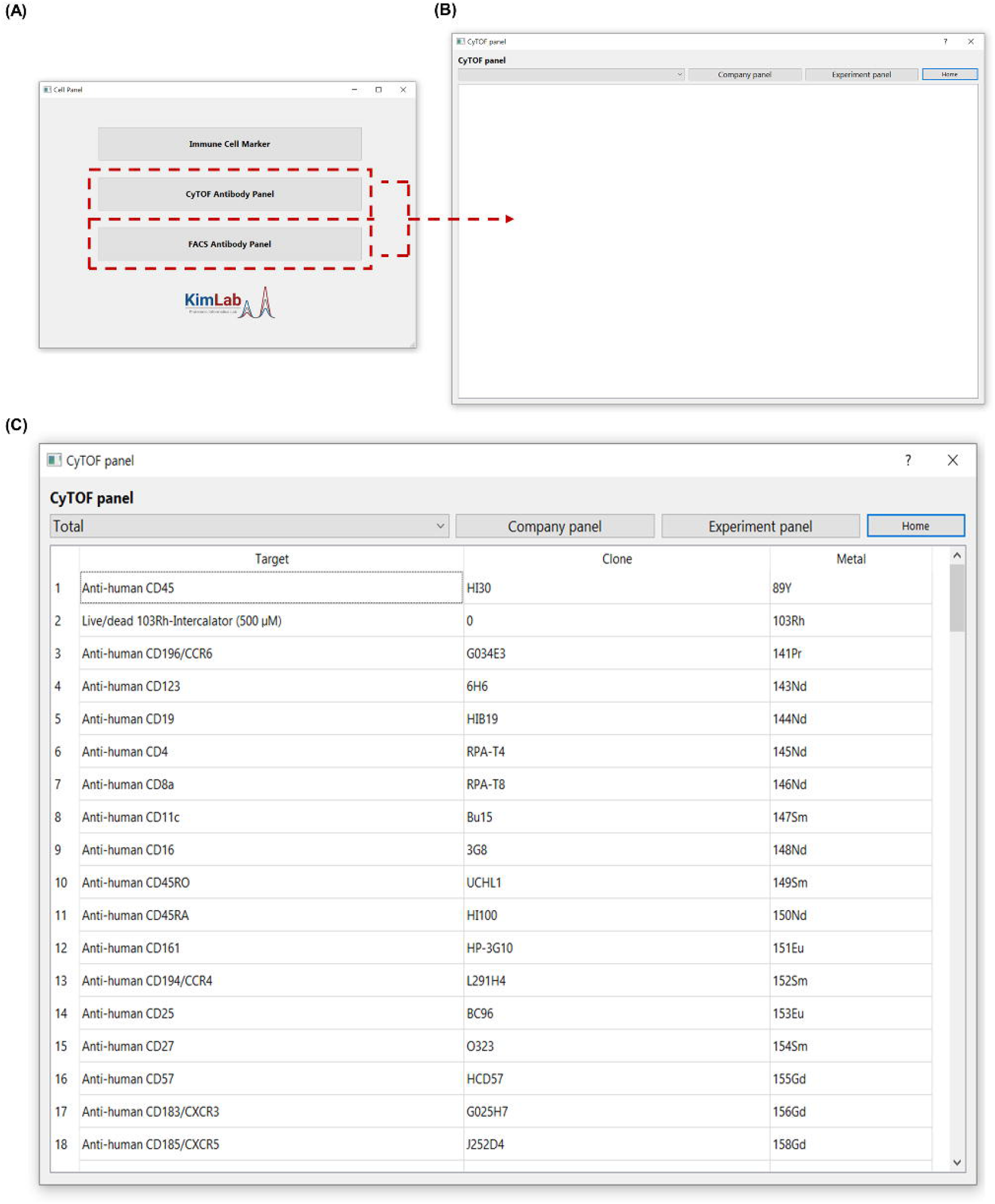
Antibody panel selection interface for mass cytometry applications in ImapDB. (A) Main window showing access points for cytometry-specific panel databases. (B) Navigation to the CyTOF Antibody Panel Dialog interface. (C) Panel selection interface featuring dual access modes for commercial and experimental panels. The example demonstrates retrieval of a commercial CyTOF panel with complete antibody specifications displayed in the table widget. Panel information includes marker targets, metal conjugates, and source references accessible through integrated DOI links.

### Data Curation and Sources

Immune cell marker information was systematically compiled from commercial immunology guidebooks (Thermo Fisher, BD Biosciences, Invitrogen) and peer-reviewed literature [12-36]. Markers are categorized by expression pattern (surface/intracellular) and cellular lineage, with secreted markers excluded from the current version (**Table 1, Supplementary Table 1**).

Antibody panel data encompasses both commercial products and custom research configurations documented in publications [37-58]. Each panel entry includes complete marker composition, clone information where available, and source documentation through catalog numbers or digital object identifiers (**Table 2, Supplementary Table 2**). This dual approach ensures coverage of standardized commercial solutions alongside innovative custom panels developed for specialized applications.

## Results

### Comprehensive Database of Human Immune Cell Markers

ImapDB provides curated marker information for human immune cell populations in their normal physiological state, encompassing both lymphoid and myeloid lineages **(Figure 1)**. The lymphoid compartment includes detailed profiles for T cells, B cells, and NK cells, with further subdivision into functionally distinct subsets. T cell populations span naive, memory, effector, and regulatory subsets, each annotated with lineage-defining and activation-associated markers. B cell categories cover developmental stages from progenitors to plasma cells, while NK cell subsets include both CD56bright and CD56dim populations. The myeloid compartment comprises monocyte subsets (classical, intermediate, and non-classical), dendritic cell populations (conventional and plasmacytoid), and granulocyte lineages including neutrophils, eosinophils, basophils, and mast cells **(Table 1)**.

Each cell population entry provides comprehensive marker information organized in tabular format for rapid reference. Surface markers accessible for live cell analysis are distinguished from intracellular markers requiring fixation and permeabilization. This systematic organization enables researchers to identify appropriate marker combinations for their specific experimental objectives while understanding the technical requirements for detection **(Supplementary Table 1)**.

### Curated Repository of Antibody Panels

The antibody panel database integrates commercial products with custom research configurations, providing researchers with diverse options for experimental design. Commercial panels from major vendors offer validated solutions for common applications, ranging from basic lineage identification to comprehensive immune profiling. These standardized panels have undergone rigorous quality control and provide consistent performance across different sample types. Custom research panels extracted from peer-reviewed publications demonstrate innovative approaches to specialized biological questions, often incorporating novel marker combinations or optimized protocols for challenging applications (**Table 2**).

Panel documentation includes complete antibody specifications with clone information, conjugate details, and recommended concentrations where available. Each entry links to source documentation through catalog numbers for commercial panels or DOIs for published protocols, enabling researchers to access detailed methodology and validation data. This comprehensive approach facilitates both direct adoption of existing panels and informed modification for specific research needs (**Supplementary Table 2**).

### User Interface and Functional Features

The ImapDB interface implements intuitive navigation through hierarchical menus that mirror conventional gating strategies used in cytometry analysis (**Figure 2**). Users searching for specific cell markers proceed through sequential selection of lineage, cell type, and subpopulation, with each level dynamically updating based on previous selections. For example, identifying markers for regulatory CD8+ T cells involves selecting Lymphocyte → CD8+ T cell → Regulatory T cell (Treg), after which relevant markers are displayed in tabular format.

Antibody panel searches utilize separate interfaces for CyTOF and FACS applications, acknowledging the distinct technical requirements of each platform (**Figure 3**). Within each interface, users can toggle between commercial and experimental panels through clearly labeled buttons. The “Company panel” option provides access to vendor-validated products, while “Experimental panel” reveals custom configurations from research literature. Selection of “Total” from the combo box displays all available panels, enabling comprehensive comparison of different approaches. The table widget presents complete panel information including all markers, conjugates, and source references, with DOI links providing direct access to primary publications.

## Discussion

ImapDB addresses a fundamental challenge in modern immunology by bridging the gap between theoretical knowledge of immune markers and practical implementation of cytometry experiments. The proliferation of single-cell technologies has revealed unprecedented cellular heterogeneity, yet this complexity creates barriers to experimental design, particularly for researchers entering the field or expanding into new biological systems. By providing an integrated platform that combines marker information with validated antibody panels, ImapDB democratizes access to advanced immunophenotyping techniques while promoting standardization across laboratories.

The hierarchical organization of the database reflects careful consideration of how immunologists conceptualize cellular relationships and design experiments. Rather than presenting markers as isolated entities, ImapDB preserves biological context through its navigation structure, facilitating both education and practical application. The distinction between commercial and custom panels acknowledges that different research contexts require different solutions—standardized panels for clinical studies demanding reproducibility, and innovative custom panels for exploratory research pushing methodological boundaries.

A key strength of ImapDB lies in its curation approach, which combines authoritative commercial resources with peer-reviewed research literature. This dual strategy ensures both breadth of coverage and depth of innovation. Commercial sources provide standardized definitions widely accepted in the field, while research publications contribute cutting-edge approaches that may not yet be commercially available. The requirement for literature support maintains scientific rigor while providing researchers with primary sources for detailed protocols beyond what can be captured in database fields.

The current version of ImapDB represents a foundation for continued development and expansion. While focusing on normal human immune cells provides essential baseline information, we recognize that disease states often alter marker expression patterns significantly. Future iterations will incorporate disease-specific profiles, beginning with well-characterized conditions where extensive phenotyping data exists. Additionally, integration with emerging technologies such as spectral flow cytometry and single-cell RNA sequencing will enhance the database’s utility as these platforms become more prevalent.

Technical sustainability relies on the open-source distribution model, which enables community contributions while maintaining quality through version control and curation standards. Regular updates through GitHub ensure the database remains current with evolving immunological knowledge, while user feedback drives feature development and content expansion. The familiar Excel-like interface and standard file formats facilitate adoption without requiring specialized training, removing barriers to access for the global research community.

The impact of ImapDB extends beyond individual experiments to influence broader research practices. By promoting the use of validated panels and standardized marker sets, the database enhances reproducibility—a critical concern in immunological research. The educational value for trainees learning cytometry techniques cannot be overstated, as exposure to diverse panel designs accelerates understanding of both technical principles and biological applications. Furthermore, the aggregation of panel information may reveal patterns in marker usage that inform future reagent development and technology advancement.

In conclusion, ImapDB provides essential infrastructure for modern immunological research by consolidating fragmented resources into an accessible, comprehensive platform. The database facilitates experimental design, promotes standardization, and accelerates research progress by eliminating redundant optimization efforts. As the complexity of immune profiling continues to increase, resources like ImapDB become increasingly vital for translating technological capabilities into biological insights. Through continued development and community engagement, ImapDB will evolve to meet emerging challenges while maintaining its core mission of democratizing access to advanced immunophenotyping techniques.

## Data Availability

All data and software are freely available at https://github.com/kimlab-cnu/Cell_Marker_GUI under MIT license. The repository includes complete database files, source code, documentation, and installation guides.

## Acknowledgements

This work was supported by the National Research Foundation of Korea (NRF) grant funded by the Korean government (MSIT) (RS-2023-00209456) and the Korea Basic Science Institute (National Research Facilities and Equipment Center) grant funded by the Korean government (MSIT) (RS-2024-00402298). We acknowledge the use of Claude Opus 4.1 solely for linguistic refinement and grammatical corrections in manuscript preparation. All scientific content, data analysis, and intellectual contributions presented herein were developed independently by the authors without the use of generative AI tools. Figure 1 was created with BioRender.com.

## Author Contributions

S.C., J.J.: conceptualization, data analysis, visualization; E.H.: data analysis; A.S., H.K.: conceptualization, methodology, project administration, funding acquisition, supervision, writing-review & editing. All authors reviewed and approved the final manuscript.

## Tables

Table 1. Summary of immune cell populations and marker categories in ImapDB.

Table 2. Overview of antibody panels included in the database.

## Supplementary Materials

Supplementary Table 1. Complete immune cell marker database with source references.

Supplementary Table 2. Detailed antibody panel specifications with validation information.

